# Genomic and phylogenomic insights into the family *Streptomycetaceae* lead to proposal of *Charcoactinosporaceae* fam. nov. and 8 novel genera with emended descriptions of *Streptomyces calvus*

**DOI:** 10.1101/2020.07.08.193797

**Authors:** Munusamy Madhaiyan, Venkatakrishnan Sivaraj Saravanan, Wah-Seng See-Too

**Affiliations:** Temasek Life Sciences Laboratory, 1 Research Link, National University of Singapore, Singapore 117604; Department of Microbiology, Indira Gandhi College of Arts and Science, Kathirkamam 605009, Pondicherry, India; Division of Genetics and Molecular Biology, Institute of Biological Sciences, Faculty of Science, University of Malaya, Kuala Lumpur, Malaysia

**Author notes:** **Corresponding author:** Temasek Life Sciences Laboratory, 1 Research Link, National University of Singapore, Singapore 117604. All these authors have contributed equally to this work.

**Keywords:** *Actinobacteria*, genome-based phylogeny, genome indices, *Actinomycetes*, Genome based classification

## Abstract

*Streptomycetaceae* is one of the oldest families within phylum *Actinobacteria* and it is large and diverse in terms of number of described taxa. The members of the family are known for their ability to produce medically important secondary metabolites and antibiotics. In this study, strains showing low 16S rRNA gene similarity (<97.3 %) with other members of *Streptomycetaceae* were identified and subjected to phylogenomic analysis using 33 orthologous gene clusters (OGC) for accurate taxonomic reassignment resulted in identification of eight distinct and deeply branching clades, further average amino acid identity (AAI) analysis showed lower AAI values or AAI within the range of 60-80 % which was previously observed in related but different genera of bacteria. The whole genome phylogeny based on concatenated core genes and AAI analyses supported the claim that those phylogenetically distinct members may be assigned to 8 novel genera namely *Actinoacidiphila, Actinomesophilus, Charcoactinospora, Curviacidiphilus, Kafeoacidiphilus, Mangroviactinospora, Peterkaempfera,* and *Streptantibioticus.* In addition, based on the core genome phylogeny and 16S rRNA tree topology and distinct chemotaxonomic and physiological properties, the sequence belonged to *Streptomyces thermoautotrophicus* was assigned to a novel genera *Charcoactinospora* which is placed under novel family *Charcoactinosporaceae*. Lastly, a clade comprising of strains that showed high 16S rRNA gene similarity (100 %) with similar tree topology in phylogenetic trees was subjected to overall genome related indices analyses such as digital DNA – DNA hybridization, and average nucleotide identity that supported the claim that *Streptomyces asterosporus* is a later heterotypic synonym of *Streptomyces calvus.*

## Introduction

The family *Streptomycetaceae* contains large numbers of highly diverse filamentous *Actinobacteria* with 895 child taxa including validly published name and synonyms. The family was first described by Waksman and Henrici [1] and was recently placed within the class *Actinomycetia* [2] and order *Actinomycetales* [3]. Members of this family are Gram positive, aerobic, non-acid-alcohol-fast organisms, mainly due to absence of mycolic acid and forms extensively branched substrate mycelium, with hypha generally non-septate and rarely fragments into conidia [4, 5]. Initially, the taxonomic assignment of first novel genus within the family *Streptomycetaceae* was based on their morphological characteristics [5] and as stated in the 8^th^ edition of Bergey’s Manual of Determinative Bacteriology, initially only four genera including *Microellobospora, Sporichthya, Streptomyces,* and *Streptovirticillium* were grouped into this family [6]. Other important genera latter added to *Streptomycetaceae* includes Genus *Kitasatospora* with *K. setae* as the type species, it was later reclassified into genus *Streptomyces* [7, 8]. However, the genus *Kitasatospora* was revived based on the (i) difference in their *meso-* A2pm to LL-DAP in the whole-cell hydrolysates, and (ii) galactose was identified as cell wall sugar only in *Kitasatospora,* as compared with *Streptomyces* [9]. Another genus proposed as a member to this family is *Streptacidiphilus,* whose members were isolated from acid soils and litter [10, 11]. Another proposed genus of this family was *Allostreptomyces* which comprises of two species [12, 13]. A genome-based taxonomic classification study has addressed the hierarchical affiliation problems in *Streptomycetaceae,* the finding has also led to demarcation of two novel genera namely *Embleya* and *Yinghuangia* [14]. Currently, this family comprised of six genera including *Allostreptomyces, Embleya, Kitasatospora, Streptacidiphilus, Streptomyces* and *Yinghuangia* [1, 7, 10, 12].

Initially, taxonomic classification of *Streptomycetaceae* was mainly based on the morphological characteristics [1] which lasted for several decades. However, several studies clearly showed that besides the colony morphology, most of the taxonomic markers including 16S rRNA gene were unable to provide a well-resolved phylogeny within the family *Streptomycetaceae* [15, 16]. The study by Labeda *et al* [17] showed the importance of the phylogenetic classification using the 16S rRNA gene sequences, it clearly illustrated the species diversity with *Streptomycetaceae,* in the form of 130 statistically supported clades and several unsupported and single member clusters. However, the phylogenetic resolution of the taxa was insufficient when 16S rRNA gene sequences alone was used for typing the strains. In addition, techniques like multilocus sequence typing (MLST) using five housekeeping genes has provided a better taxonomic resolution based on the evolutionary relationships among the species of *Streptomyces.* This MLST analysis also supported the phylogenetic distinctness of other closely related genera such as *Kitasatospora* and *Streptacidiphilus.* Interestingly, this study also supported the transfer of several *Streptomyces* species to the genus *Kitasatospora* and also reducing 31 species clusters to a single taxon [18].

Alternatively, genome-based studies are emerging as a promising approach for delineation of genera and species within *Streptomycetaceae* [14, 16, 19]. Difference in the similarity or distance between the genomes are attributed to Overall Genome Related Index (OGRI) that comprises of dDDH (digital DNA-DNA hybridization), ANI (average nucleotide identity), AAI (average amino acid identity) that are used for genus, species and subspecies delineation [20]. In particular for higher taxonomic placement at genus and family levels, orthologous gene clusters (OGCs) were concatenated and constructed as core-genome based phylogenomic trees similar to a multiple-locus sequence typing performed with a few housekeeping genes. Thus, the combination of genomic data with highly conserved phenotypic traits and chemotaxonomic markers have proved to be helpful for taxonomic assignment in higher ranks [20]. In a previous study, different taxa of *Actinobacteria* were subjected to genome level analyses which resulted in identification of 2 orders, 10 families, 17 genera and transfer of more than 100 species to other genera. This study also suggested that genome size and DNA G+C content were to be considered as important taxonomic markers in *Actinobacteria* taxonomy [14]. The present study was designed to elucidate genome-based evolutionary relationships between the taxa of *Streptomycetaceae,* currently available in the database (https://lpsn.dsmz.de/family/streptomycetaceae) for assignment of correct taxonomic affiliation. Notably, a recent OGRI analyses on the available genomes of *Streptomyces* genus has identified 6 clades that comprised of strains that were reclassified as heterotypic synonym [21].

## Materials and methods

### 16S rRNA and OGC based core genome analysis

To understand the evolutionary relationship between the taxa of *Streptomycetaceae,* 16S rRNA gene sequences of 216 type strains (Supplementary Table S1) that with genomic data were retrieved from the EzBioCloud Database [22] on 6 February, 2020. The CLUSTALW tool of MEGA version 7 was used for multiple sequence alignments, pairwise comparison and phylogenetic analysis of the individual sequences [23]. Distances were calculated using the Kimura correction in a pairwise deletion [24]. Using MEGA 7 software with 500 non-parametric bootstrap replicates, a neighbor-joining based phylogenetic tree was constructed.

To ascertain the result of 16S rRNA gene analysis, whole genome-based phylogeny analysis was carried out for various taxa of *Streptomycetaceae.* From a total of 228 strains available with whole genome sequence, 12 strains were not included due to fragmented genomic data and low quality of sequences. Therefore, 216 taxa of the *Streptomycetaceae* were retrieved from NCBI (https://www.ncbi.nlm.nih.gov/) on 6 February, 2020 (Supplementary Table S1). The identities of the genomes were verified using the 16S rRNA gene sequences in the EzBioCloud database [22]. OGCs were then identified using the panX pan-genome pipeline. A total of 33 core genes (Supplementary Table S2) were identified and were then subjected to concatenated alignment [25]. With the concatenated alignment built from these OGCs, suitable evolutionary model for each OGC was determined using Modeltest-NG [26]. Finally, the concatenated OGCs were subjected to maximum likelihood based phylogenetic analysis using RAxML-NG [27].

### Analysis of OGRI indices

The complete genome-based comparison in the form of OGRI analysis such as digital DNA-DNA hybridization (dDDH), average nucleotide identity (ANI) and average amino acid identity (AAI) were calculated for strains used in this study. Genome distance in the form of dDDH values were calculated by the Genome-to-Genome Distance Calculator (version 2.1; http://ggdc.dsmz.de) [28]. The ANI was calculated based on the blastn method for close relatives using EDGAR v2.3 (https://edgar.computational.bio.uni-giessen.de) [29]. The pairwise average AAI was calculated using the AAI workflow with default settings in CompareM v0.0.23 (https://github.com/dparks1134/CompareM).

## Results and discussion

The 16S rRNA gene and phylogenomic analyses have revealed the presence of eight distinct phylogenetic lineages, which exhibit low 16S rRNA gene similarity (<97.3 %) with the other type strains of *Streptomycetaceae* and unique tree topology with high bootstrap support within their clade members (Fig. 1 and Fig. S1). For instance, *Streptacidiphilus griseoplanus,* a reclassified *Streptomyces* and *Streptacidiphilus bronchialis,* a ciprofloxacin resistant bacterium isolated from human clinical specimen produced different coloured aerial mycelium while *S. griseoplanus* produced grey mycelium, while their 16S rRNA gene sequence was subjected for phylogenetic analysis, both shared 98.82 % similarity with high bootstrap support. The two species also recorded an AAI value of 69.05 and 69.16 % with the *Streptacidiphilus* type species *S. albus* (Fig. 2 and Table S3). Coherent grouping of this taxa was evident through visualization of the AAI data as heat maps (Fig. 2). Distinct tree topology recovered in both 16S rRNA gene sequence analysis and core gene phylogeny combined with the low AAI values recorded with the type species of *Streptacidiphilus* strongly supported the proposal of a novel genus *Peterkaempfera* accommodating *Streptacidiphilus griseoplanus* and *Streptacidiphilus bronchialis* strains. Members of this genera possessed galactose, ribose and traces of mannose in the whole cell hydrolysates with MK9 (H6) as hallmark menaquinone (Table 1).

**Fig. 1.**
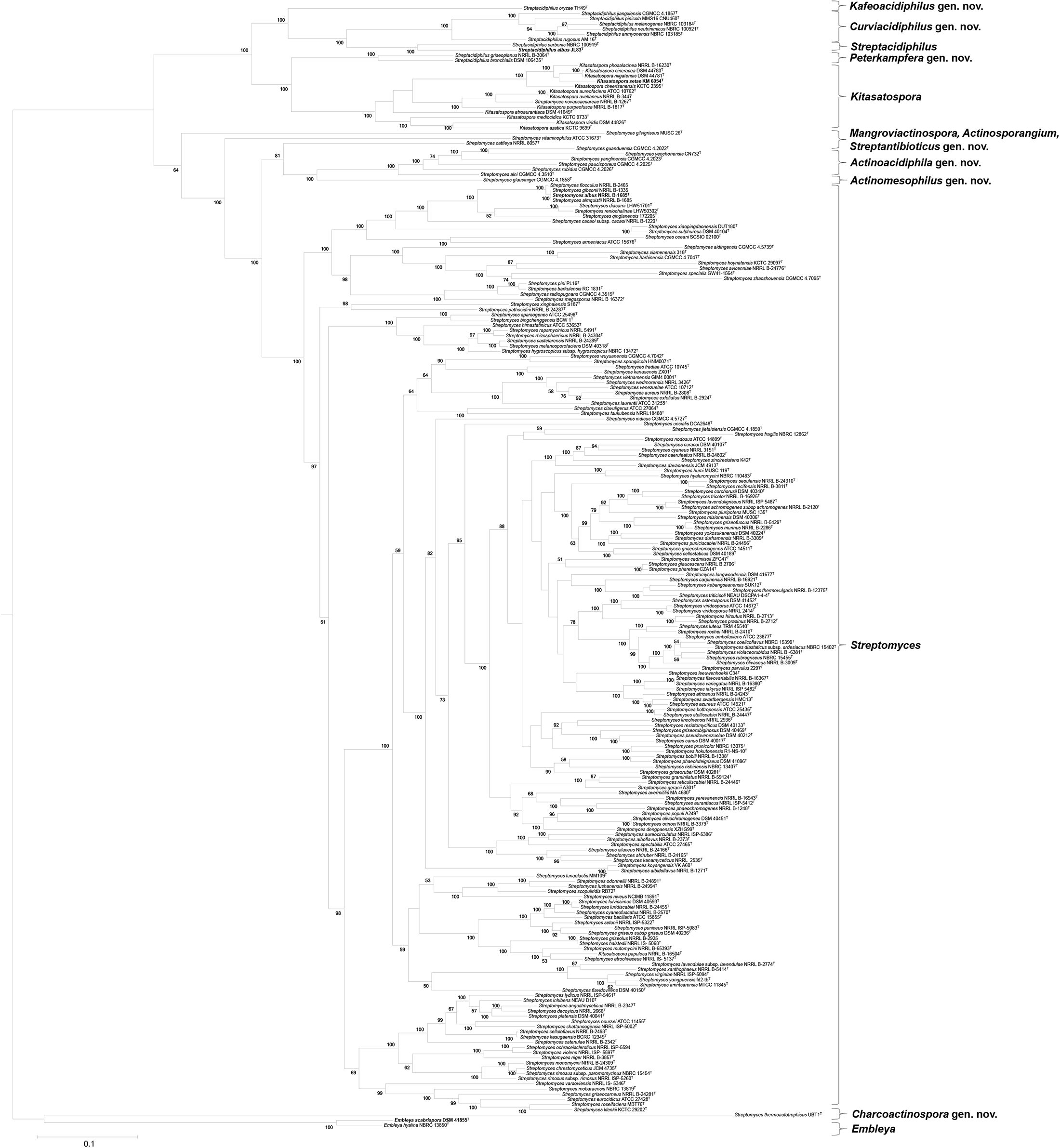
Maximum likelihood tree of core genome-based phylogeny of *Streptomyces* species. Tree generated using a core genome alignment (33 genes, 15,708 bps) with RAxML-NG. Bootstrap values represent the percentage of concurring bootstraps from 100 iterations. Scale bar, 0.10 nt substitutions per position. Species, strains name and accession numbers with assembly levels are listed in Table S1.

**Table 1.**
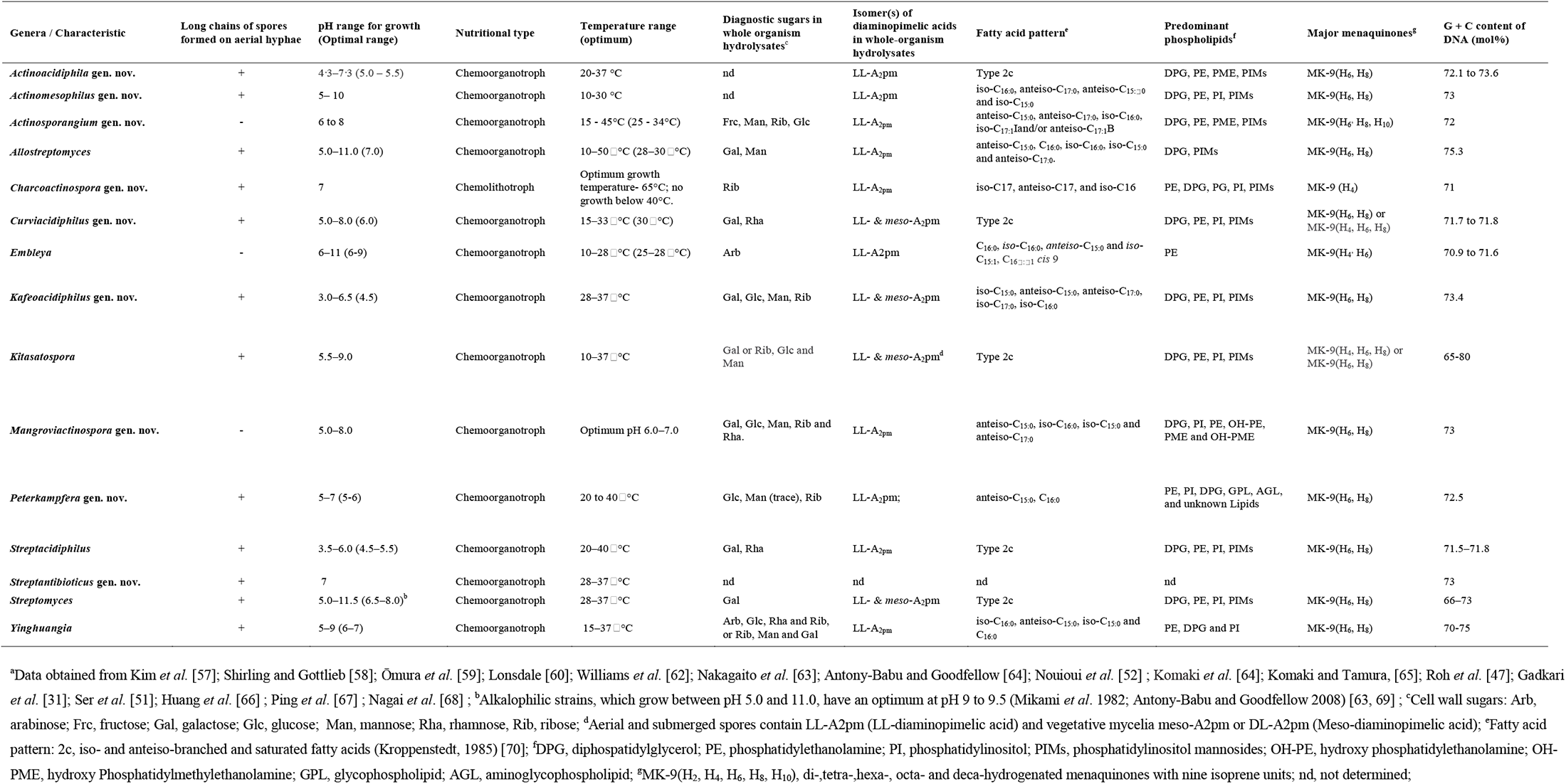
Morphological, physiological, and chemotaxonomic characteristics of currently available and newly proposed genera of *Streptomycetaceae* family^a^

Another clade of *Streptacidiphilus* comprised of 6 type strains *(S. jiangxiensis, S. pinicola, S. melanogenes, S. neutrinimicus, S. anmyonensis* and *S. rugosus)* shared a low 16S rRNA gene similarity (96.67-98.75 %), while their coherent clade formation was reflected in the core genome based phylogenetic analysis (Fig. 1), those members showed an AAI value ranged from 71.01 – 71.86 % on comparison with the *Streptacidiphilus* type species *S. albus* (Fig. 2 and Table S3). When their AAI data sets were visualized as a heat map, all these 6 strains coherently group into distinct taxa (Fig. 2). Distinct tree topology recovered in both 16S rRNA gene sequence analysis and core gene phylogeny combined with the low AAI values supported the proposal of *Curviacidiphilus* genus nov. to accommodate those six *Streptacidiphilus* strains (*S. jiangxiensis, S. pinicola, S. melanogenes, S. neutrinimicus, S. anmyonensis* and *S. rugosus)* among the chemotaxonomic markers galactose and rhamnose were identified as diagnostic sugars in the whole cell hydrolysates of the organism and cell wall contained LL- & *meso*-A_2_pm (Table 1).

**Fig. 2.**
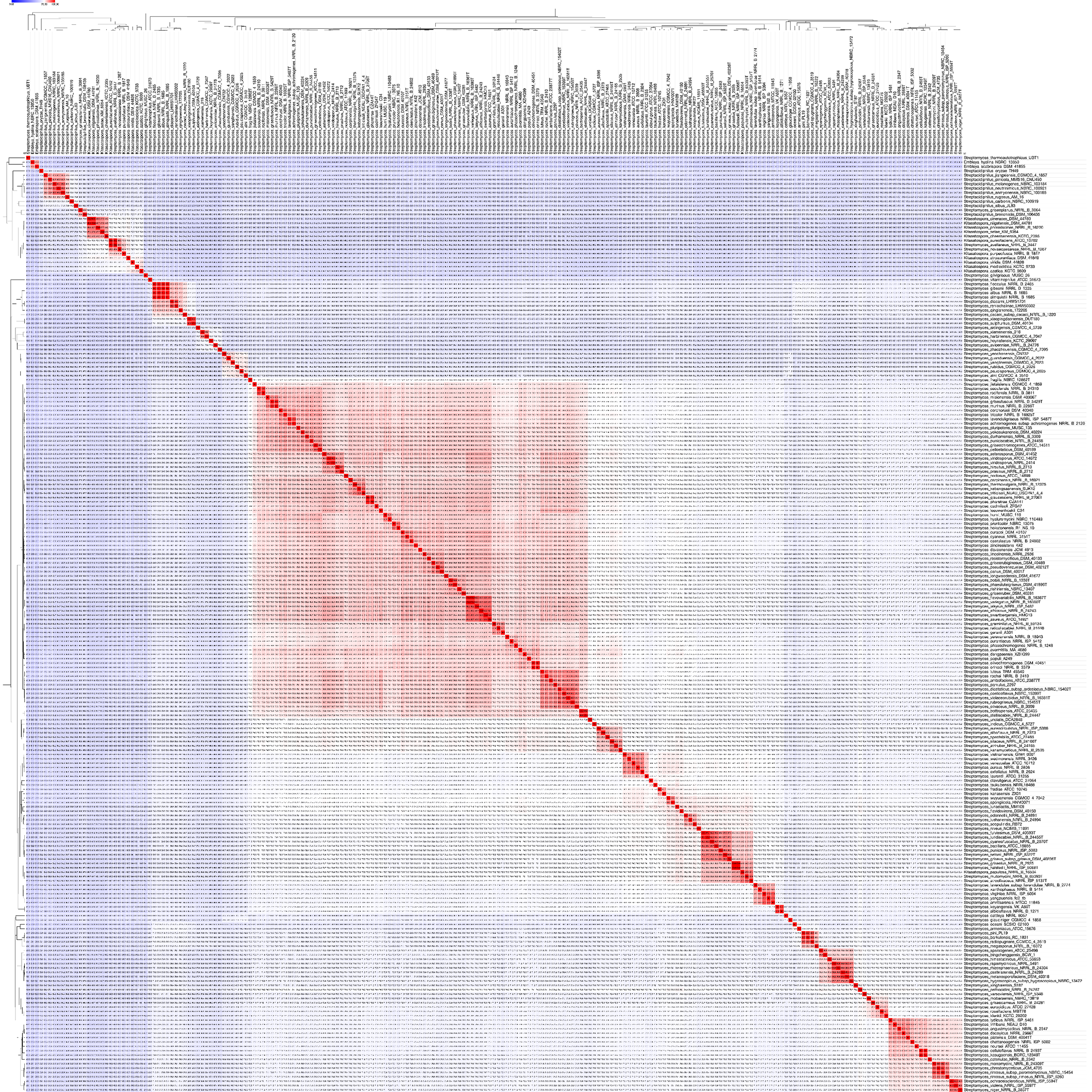
AAI from pairwise whole-genome comparisons within the family *Streptomycetaceae*. The heat map shows AAI values between genomes, along with the tree cladogram to show relationships. Boxed regions indicate inferred genus clusters with at least two members based on AAI comparisons, as well as monophyly in the genome-based phylogeny. Species, strains name and accession numbers with assembly levels are listed in Table S1.

*Streptacidiphilus oryzae* belonged to acidophilic group of actinomycetes recovered from the acidic soils of rice paddies, the bacterium grows between pH 3.0 to 6.5. When their 16S rRNA gene was analyzed phylogenetically closest member was *S. griseoplanus* with 97.36 % similarity. Similarly, single species containing clade comprising of *Streptacidiphilus oryzae* recorded an AAI range of 67.5-68.8 % with other *Streptacidiphilus* species. The AAI value of *S. oryzae* with *S. griseoplanus* is 68.41% (Fig. 2 and Table S3). Distinct tree topology recovered in both 16S rRNA gene sequence analysis and core gene phylogeny combined with the low AAI values supported the proposal of novel genus *Kafeoacidiphilus* to accommodate *S. oryzae,* apart from *Streptacidiphilus,* this is the only other genera within *Streptomycetaceae* whose member has the ability to grow at a low pH 3.0 – 6.5, the diagnostic sugar in the whole cell hydrolysates includes galactose, glucose, mannose and ribose, the cell wall contained LL- & *meso*-A_2_pm (Table 1).

Presence of distantly evolved but unique lineages were also identified in the genus *Streptomyces.* A lineage representing a clade comprising of *S. alni, S. guanduensis, S. paucisporeus, S. rubidus, S. yanglinensis,* and *S. yeochonensis* shared low 16S rRNA gene similarity range of 97.3–96.1 % in comparison with *S. albus,* the type strain of *Streptomyces.* In addition, members of this clade showed low 16S rRNA gene similarity range (97.37-96.33 %) to *S. cattleya,* a single species cluster that is phylogenetically closest to these six taxa. Owing to low 16S rRNA gene similarity percent with other type strains and deeply branching in tree topology of this clade in both 16S rRNA gene sequence (Fig. 1) and OGCs based core gene analysis, it was subjected to AAI analysis. The members recorded a wide AAI range of 61.75-79.36 % (mean: 70.08 %) with the type species of the *Streptomyces,* all these six strains together form into a distinct clade which was reflected in the AAI data-based heat map (Fig. 2 and Table S3). These values are within the range 60-80 %, which was previously observed in a genomic and metagenomic taxonomic classification of related but different genera [30] and are within the range recorded for already established genera of *Streptomycetaceae.* For instance, when the type strain of *Kitasatospora setae* was compared with other type species of *Streptomyces,* AAI in the range of 61.62–73.79 % (mean – 67.7 %) was recorded (Fig. 2 and Table S3). Distinct tree topology recovered in both 16S rRNA gene sequence analysis and core gene phylogeny combined with the low AAI values with the type species of *Streptomyces* supported the proposal of *Actinoacidiphila* genus nov. to accommodate *S. alni, S. guanduensis, S. paucisporeus, S. rubidus, S. yanglinensis,* and *S. yeochonensis*.

Single cluster lineages such as *S. gilvigriseus, S. cattleya* and *S. glauciniger* showed lowest 16S rRNA gene sequence similarity of 96.2, 97.65 and 96.53 % with their nearest taxa namely *S. quinglanensis, S. chrestomyceticus* and *S. alini* respectively. Similarly, *S. gilvigriseus* recorded ANI, dDDH and AAI values of 74.83, 20.8 and 64.05 % with *S. quinglanensis* respectively. *S. cattleya* recorded ANI, dDDH and AAI values of 77.77, 22.4 and 70.38 % with *S. chrestomyceticus* respectively. *S. glauciniger* showed ANI, dDDH and AAI values of 78.25, 22.6 and 72.53 % with *S. alni* respectively. Distinct tree topology recovered in both 16S rRNA gene sequence analysis and core gene phylogeny combined with the low AAI values supported the proposal of three novel genera namely *Mangroviactinospora, Streptantibioticus* and *Actinomesophilus* to accommodate *S. gilvigriseus, S. cattleya* and *S. glauciniger* respectively.

Among the different clusters analysed in this study, *S. thermoautotrophicus* is phylogenetically distinct and forms deep clade among those branches within the family *Streptomycetaceae*. This strain is a Gram-positive thermophile originally isolated from soil covering burning charcoal pile and it is an obligate chemolithotrophic organism using CO and CO_2_ plus H_2_ as substrates, with preferable thermophilic temperature (65 °C) for growth [31]. At the time of its description, it was placed within the genus *Streptomyces* based on its DNA G+C content (71 mol %), phospholipid pattern type-II and MK-9 (H4) as the major quinone (refer Table 1). This clade has an AAI value range of 60.89 – 63.65 % with other *Streptomyces* species, and AAI value range 62.7-62.81 % with *Embleya* species (Fig. 2 and Table S3), all these polyphasic traits strongly suggest to propose is as novel genera, such a proposal was already hinted in an earlier study [32]. One of the closest neighbour identified in 16S rRNA gene phylogenetic and the phylogenomic analyses. *Embleya scabrispora* DSM 41855^T^ is the closest taxon of *S. thermoautotrophicus* within family *Streptomycetaceae* recording 91.85 % 16S rRNA gene similarity, other OGRI’s are 74.07 and 20.6 % of ANI and dDDH respectively. The AAI values recorded were within the range (60-80 %) previously observed in a study for phylogenetically related but different genera [30]. The 16S rRNA and phylogenomics based tree topology shows that this sequence belongs to a novel clade that was evolutionarily distant from the nearer clades (Fig. 1 and Fig. S1). For instance, 16S rRNA based neighbour joining tree shows that *Motilibacter rhizosphaerae* and *Cryptosporangium aurantiacum* are closest organisms but belonging to different families such as *Motilibacteraceae* and *Cryptosporangiaceae* respectively, both strains showing 93.8 % similarity with of *S. thermoautotrophicus.* Another strain that is phylogenetically closer to *S. thermoautotrophicus* is *Acidothermus cellulotlyticus* with 94.71 % similarity, which also belonged to a unique family *Acidothermaceae* (Fig. S2). Thus with a genetic distance of 94.71 – 93.8 % 16S rRNA gene similarities within the inter genus of *Motilibacter, Cryptosporangium* and *Acidothermus* novel family were previously proposed [33, 34, 35], in line with this observation in the present study, a novel family *Charcomycetaceae* is proposed to accommodate *S. thermoautotrophicus* that showed 16S rRNA gene sequence similarity of 91.85 % with *Embleya scabrispora* DSM 41855^T^, a genetically closer taxon of *Streptomycetaceae.* The major characteristics that differentiate *Charcomycetaceae* member from the *Streptomycetaceae* are presence of ribose as diagnostic sugar in the whole cell hydrolysates of the organism, the predominant menaquinone was MK-9 (H4) (Table 1). The ability of the strain to grow in a thermophilic temperature (65 °C) and no growth below 40 °C, chemolithotrophic growth on CO, H_2_ plus CO_2_ and unable to grow in a complex medium containing heterotrophic substrate such as sugar, organic acid, alcohols and amino acids (Table 1).

A cluster containing *Streptomyces vitaminophilus* was originally named as *Actinosporangium vitaminophilium* [36]. Based on physiological properties and certain chemotaxonomic characteristics (cell wall type, type-II phospholipid pattern, presence of hexa and octa hydrogenated menaquinones), it was transferred to genus *Streptomyces* [37]. However, in the present study, this strain was observed to form a deeply branching clade on 16S rRNA gene and core genome-based phylogenomics trees. In addition, the AAI values of phylogenetically closest members of the core genome tree namely *S. cattleya* and *S. gilvigriseus* are 69.4 and 65.4 % respectively (Fig. 2 and Table S3), suggested that the genus *Actinosporangium* need to be revive to accommodate the members of this clade as its AAI values with closest neighbour were well within the range 60-80 % observed on phylogenetically closer but related genera [30].

Interestingly, the five single-species containing clusters and three multiple-species containing clades on subjected to 16S rRNA gene sequence and core genome phylogeny analysis revealed that their type strains are equally distant from the other *Streptomyces* members as a *Kitasatospora* or *Embleya* taxa is from *Streptomyces.* Previous studies also showed the distinctness of the genera status for *Kitasatospora, Streptomyces* and *Streptacidiphilus* through 16S rRNA gene sequence and MLST analyses [17, 18].

Based on the low 16S rRNA gene similarity and tree topology in both 16S rRNA gene and OGCs-based phylogenetic analyses, it is clearly revealed the presence of taxa within the family *Streptomycetaceae* that required reclassification at genus level. In a previous attempt to construct the genome taxonomy database, phylogeny was built with the dereplicated subset of all sequenced bacterial genomes, to determine the hierarchical ranks of phylum *Actinobacteria* that hypothesized the presence of several novel genera within *Streptomycetaceae* [38].

On comprehensive phylogenetic analysis of taxa within *Streptomycetaceae,* a clade comprising of two different species (*S. asterosporus* and *S. calvus)* shared 100 % 16S rRNA gene similarity and with high bootstrap support in phylogenetic trees. These clades were carefully analysed using various OGRI, to understand their exact taxonomical affiliation. *S. asterosporus* and *S. calvus* recorded 92.9 % and 99.12 % of dDDH and ANI, respectively (Table 2). These values were above the threshold level for species assignment, the proposed cutoff values of the OGRIs dDDH, ANI and AAI are > 70 %, 95-96 % and >95 %, respectively [39, 40, 41]. The genomic evidence supports that one strain in this clade may be a heterotypic synonym. In addition to similarities in OGRI, physiological and biochemical characteristics between the strains in each of these two clades are presented in the Supplementary Tables S4. Most of the biochemical traits are similar among the taxa of this clade, only a little difference was observed among these strains. For instance, critical analysis of biochemical properties of *S. asterosporus* and *S. calvus* revealed most of the properties were similar for both the strains and a variation was noticed only in D-xylose and raffinose utilization. Both the genomic and biochemical properties strongly support the proposal that *Streptomyces asterosporus* may be a later heterotypic synonym of *Streptomyces calvus.*

**Table 2.** 16S rRNA gene sequence percent identity and overall genome-relatedness indices for specific clades in the *Streptomycetaceae* family.

| Type strain(s) | 16S %<br>similarity | dDDH* | ANI<sup>†</sup> |
| --- | --- | --- | --- |
| *Streptomyces asterosporus* DSM 41452<sup>T</sup> | 100 | 92.9 | 99.12 |
| *Streptomyces calvus* CECT 3271<sup>T</sup> |  |  |  |

\*dDDH similarities were calculated using Genome-to-Genome Distance Calculator (formula 2) web server 2.1 (http://ggdc.dmz.de).
<sup>†</sup>Average nucleotide identities (ANI) is a similarity measure between two genome sequences calculated using an ANI calculator (https://www.ezbiocloud.net/tools/ani).

In conclusion, apart from identifying this heterotypic synonym, genome level phylogenetic (16S rRNA gene) and phylogenomic (core genome) approaches combined with OGRI like AAI had revealed the presence of 8 novel phylogenetic lineages, whose taxa are proposed as 8 novel genera, their descriptions are provided herewith.

## Taxonomic consequences: New taxa

### Description of *Charcoactinosporaceae* fam. nov

Char.co. ac.ti.no’spo.ra’ce.ae (N.L. fem. n. *Charcoactinospora,* type genus of the family; *-aceae,* ending to denote a family; N.L. fem. pl. n. *Charcoactinosporaceae,* the *Charcoactinospora* family).

The family shares chemotaxonomic and morphological features with the *Streptomycetaceae* family including traits such as Gram positive, non-acid fast due to absence of mycolic acids, non-motile, aerobic produces relatively stable branching vegetative hyphae. The main features that differentiate it from the *Streptomycetaceae* members like *Streptomyces, Kitasatospora* and *Streptacidiphilus* are its strain ability to grow at 65 °C but no growth detected below 40 °C and lithotrophy in nutrition with unique chemolithotrophic potential to use CO_2_ or H_2_ plus CO_2_ as substrates. The hall mark menaquinone identified in member is MK-9 (H4). The genomic G+C content is around 71 %. In addition, phylogenetic analyses of genome and 16S rRNA gene sequences supported the proposal of the family nov. The description is as that for *Charcoactinospora* [31] which is the type and currently sole genus of the family. It has been separated from other families based on phylogenetic analyses of genome and 16S rRNA gene sequences.

### Description of *Actinoacidiphila* gen. nov

*Actinoacidiphila* (Gr. n. *actis,* actinos a ray; a.ci.di’phi.la. N.L. neut. n. *acidum* acid; Gr. adj.*philos* loving; N.L. fem. adj. *acidiphila* acid-loving neutrotolerant actinomycetes).

Aerobic, Gram-positive, non-motile, neutrotolerant, acidophilic streptomycete that forms branched substrate and aerial hyphae. Smooth surface spores borne in flexuous or rectiflexibiles spore chain. Growth occurs at 20-37 °C. Neutrotolerant acidophilic strains grow in a pH range of 4.5-4.5 with optimum range of 5.0 – 5.5. Cell wall contained LL-diaminopimelic acid as major component, the cell membrane phospholipid type II with diphosphatidylglycerol, phosphatidylethanolamine, phosphatidylinositol and phosphatidylinositol mannosides as major polar lipids. Hexa and octa menaquinones with nine isoprene units MK-9 (H6, H8) as the predominant menaquinones. L-arabinose, D-fructose, D-raffinose, L-rhamnose, L-sorbose, D-sucrose, D-trehalose and D-xylose are used as sole carbon source at 1 % (w/v). The genomic G+C□content range was 72.1 to 73.6□mol%. The genus has been separated from *Streptomyces* based on phylogenetic analyses of genome and 16S rRNA gene sequences. The type species is *Actinoacidiphila yeochonensis*, comb. nov.

### Description of *Actinoacidiphila alni* comb. nov

*Actinoacidiphila alni* (al’ni L. gen. fem. n. *alni,* of the alder, referring to the isolation of the type strain from *Alnus nepalensis)*

Basonym: *Streptomyces alni* (Liu *et al.* 2009) EMEND Nouioui *et al.* 2018.

The description is the same as for *Streptomyces alni* [42]. Phylogenomic analyses of the core genome provided strong evidence for assignment of this species in the novel genus *Actinoacidiphila.* The G+C content of the type strain genome is 72.1 %, its approximate genome size is 8.27 Mbp, its GenBank accession is GCA_900112845.1. The type strain is CGMCC 4.3510^T^ = D65^T^ =DSM 42036^T^ = JCM 16122^T^ =NRRL B-24611^T^).

### Description of *Actinoacidiphila guanduensis* comb. nov

*Actinoacidiphila guanduensis* (N.L. masc./fem. adj. *guanduensis,* of or belonging to Guandu, the source of the soil from which the type strain was isolated).

Basonym: *Streptomyces guanduensis* Xu *et al.* 2006.

The description is the same as for *Streptomyces guanduensis* [43]. Phylogenomic analyses of the core genome provided strong evidence for assignment of this species in the novel genus *Actinoacidiphila.* The G+C content of the type strain genome is 73.1 %, its approximate genome size is 8.22 Mbp, its GenBank accession is GCA_900103985.1. The type strain is 701^T^ (= CGMCC 4.2022^T^ = DSM 41944^T^ = JCM 13274^T^ = NBRC 102070^T^).

### Description of *Actinoacidiphila paucisporeus* comb. nov

*Actinoacidiphila paucisporeus* (L. adj.*paucus,* few; N.L. adj. *sporeus,* spored; N.L. masc. adj.*paucisporeus,* few spored, forming few spores).

Basonym: *Streptomyces paucisporeus* (Xu *et al.* 2006) EMEND Nouioui *et al.* 2018.

The description is the same as for *Streptomyces paucisporeus* [43]. Phylogenomic analyses of the core genome provided strong evidence for assignment of this species in the novel genus *Actinoacidiphila.* The G+C content of the type strain genome is 72.2 %, its approximate genome size is 8.16 Mbp, its GenBank accession is GCA_900142575.1. The type strain is 1413^T^ (= CGMCC 4.2025^T^ = DSM 41946^T^ = JCM 13276^T^ = NBRC 102072^T^).

### Description of *Actinoacidiphila rubidus* comb. nov

*Actinoacidiphila rubidus* (L. masc. adj. *rubidus,* dark red).

Basonym: *Streptomyces rubidus* (Xu *et al.* 2006) EMEND Nouioui *et al.* 2018.

The description is the same as for *Streptomyces rubidus* [43]. Phylogenomic analyses of the core genome provided strong evidence for assignment of this species in the novel genus *Actinoacidiphila.* The G+C content of the type strain genome is 72.9 %, its approximate genome size is 9.01 Mbp, its GenBank accession is GCA_900110255.1. The type strain is 13c15^T^ (= CGMCC 4.2026 ^T^ = DSM 41947 ^T^ = JCM 13277 ^T^ = NBRC 102073^T^).

### Description of *Actinoacidiphila yanglinensis* comb. nov

*Actinoacidiphila yanglinensis* (N.L. masc./fem. adj. *yanglinensis,* of or belonging to Yanglin, the source of the soil from which the type strain was isolated).

Basonym: *Streptomyces yanglinensis* (Xu et al. 2006) EMEND Nouioui *et al.* 2018.

The description is the same as for *Streptomyces yanglinensis* [43]. Phylogenomic analyses of the core genome provided strong evidence for assignment of this species in the novel genus

*Actinoacidiphila.* The G+C content of the type strain genome is 72.6 %, its approximate genome size is 9.59 Mbp, its GenBank accession is GCA_900107965.1. The type strain is 1307^T^ (= CGMCC 4.2023^T^ = DSM 41945^T^ = JCM 13275^T^ = NBRC 102071^T^).

### Description of *Actinoacidiphila yeochonensis* comb. nov

*Actinoacidiphila yeochonensis* (ye.o.chon.en’sis N.L. masc./fem. adj.*yeochonensis,* of Yeochon, a province in Korea, referring to the place where the organism was first isolated).

Basonym: *Streptomyces yeochonensis* (Kim *et al.* 2004) EMEND Nouioui *et al.* 2018.

The description is the same as for *Streptomyces yeochonensis* [44]. Phylogenomic analyses of the core genome provided strong evidence for assignment of this species in the novel genus *Actinoacidiphila.* The G+C content of the type strain genome is 73.6 %, its approximate genome size is 7.82 Mbp, its GenBank accession is GCA_000745345.1. The type strain is CN 732^T^ (= DSM 41868^T^ = IMSNU 50114^T^ = JCM 12366^T^ = KCTC 9926^T^ = NBRC 100782^T^ = NRRL B-24245^T^).

### Description of *Actinomesophilus* gen. nov

*Actinomesophilus* (Ac.tino.meso.phi.lus Gr. fem. n. *aktis,* aktinos, ray; Gr. adj. *mesos,* middle; N.L. adj.*philus -a -um,* friend, loving; from Gr. adj.*philos -ê -on;* ray, loves mesophilic temperature).

Aerobic, Gram-positive mesophilic actinomycete that forms an extensively branched substrate mycelium and aerial hyphae that differentiate into long spiral spore chains with 15-20 cylindrical spores per chain. Spore surface is smooth. Grows at 10-30 °C but not at 40 °C, and a pH of 5.0 – 10.0. Cell wall is of type I, phospholipid type II with major cellular fatty acids made up of iso-C_16:0_, anteiso-C_17:0_, anteiso-C_15:□0_ and iso-C_15:0_ and menaquinone MK-9 (H_6_, H_8_). Nitrate is reduced but gelatin is not liquefied. Can able to use L-arabinose, D-fructose, meso – inositol, D-mannitol, D-raffinose, L-rhamnose, D-sucrose and D-xylose at 1 % w/v as sole carbon source. The genomic G+C content is around 72 %. The genus has been separated from *Streptomyces* based on phylogenetic analyses of genome and 16S rRNA gene sequences. The type species is *Actinomesophilus glauciniger* comb. nov.

### Description of *Actinomesophilus glauciniger* comb. nov

*Actinomesophilus glauciniger* (glau’ci.ni.ger L. masc. adj. *glaucus,* greenish grey; L. masc. adj. *niger,* black; N.L. masc. adj. *glauciniger,* greenish black, referring to the colour of colony reverse on modified Bennett’s agar).

Basonym: *Streptomyces glauciniger* Huang *et al.* 2004.

The description is same as given before [45] for *Streptomyces glauciniger* with the following additions. Phylogenomic analyses provided strong evidence for assignment of this species to the novel genus *Streptomyces.* The G+C content of the type strain genome is 72.3 %, its approximate genome size is 9.81 Mbp, its GenBank accession is GCA_900188405.1. The type strain is FXJ14^T^ (=AS 4.1858^T^ = DSM 41867^T^ = JCM 12278^T^ = LMG 22082^T^ = NBRC 100913^T^).

### Description of *Actinosporangium* gen. nov

Cells stain Gram-positive and produces new antibiotics called pyrrolomycins and is characterized by nonmotile spores, mesophilic, and a type I cell wall. Pseudosporangia are spherical to oval, and sometimes irregular and are formed on aerial mycelium; there is no definite sporangial wall. Hydrolysis of starch, liquefaction of gelatin, and reduction of nitrate were positive. Peptonization and coagulation of skim milk and formation of melanoid pigment were negative. Utilizes D-glucose, D-xylose, L-rhamnose, and glycerol for growth, but not L-arabinose, D-fructose, D-mannitol, I-inositol, sucrose, or raffinose. The major menaquinones are MK-9(H_6_), MK-9(H_8_) and MK-9(H_10_). The genomic G+C content is around 72 %. The genus has been separated from *Streptomyces* based on phylogenetic analyses of genome and 16S rRNA gene sequences. The type species is *Actinosporangium vitaminophilum* comb. nov.

### Description of *Actinosporangium vitaminophilum* comb. nov

*Actinosporangium vitaminophilum* (N.L. neut. n. *vitaminum,* vitamin; N.L. neut. adj.*philum,* friend, loving; from Gr. neut. adj.*philon;* N.L. neut. adj. *vitaminophilum,* vitamin-loving).

Basonym: *Streptomyces vitaminophilum* corrig. (Shomura *et al.* 1983) Goodfellow *et al.* 1986 EMEND. Nouioui *et al.* 2018.

The description is the same as for *Streptomyces vitaminophilum* [36, 37]. Phylogenomic analyses of the core genome provided strong evidence for assignment of this species in the novel genus *Actinosporangium.* The G+C content of the type strain genome is 72 %, its approximate genome size is 6.55 Mbp, its GenBank accession is GCA_001445835.1. The type strain is SF 2080^T^ (=ATCC 31673^T^ = DSM 41686^T^ = IFO 14294^T^ = JCM 6054^T^ = NBRC 14294^T^ =NRRL B-16933^T^).

### Description of *Charcoactinospora* gen. nov

*Charcoactinospora* (N.L. fem. n. *Charco,* named for charcoal, a source for isolation; Gr. n. *actis,* actinos a ray; Gr. fem. n. *spora,* a seed and, in biology, a spore; N.L. fem. n. *Charcoactinospora,* actinomycete with spiny spores).

Aerobic, Gram positive, non-acid fast, non-motile, thermophilic obligately chemolithotrophic bacteria. The hyphae is relatively stable branching vegetative hyphae, two to eight oval spores reside on the substrate hyphae. Endospores are not formed but synnemata, sporangia or sclerotia are formed. Cell wall belongs to type-I with LL-diaminopimelic acid and ribose. Cellular fatty acids composed of Iso and anteiso branched fatty acid pattern is synthesized, with iso-C17, anteiso-C17, and iso-C16 being the predominant fatty acids. Polar lipids comprised of phosphatidylethanolamine, diphosphatidylglycerol, phosphatidylglycerol, phosphatidylinositol, phosphatidylinositol mannosides (PL-type 2). Predominant menaquinone is MK-9 (H_4_). Chemolithotropically grows on CO, H_2_ plus CO_2_. Complex medium or heterotrophic substrates such as sugars, organic acids, amino acids and alcohols don’t support growth. Optimum growth observed at 40 °C. The genus name reflects the charcoal source used for the isolation of the type species. The description is as for *Charcoactinospora thermoautothrophicus,* comb. nov., which is the type species. The genus has been separated from *Streptomyces* based on phylogenetic analyses of genome and 16S rRNA gene sequences.

### Description of *Charcoactinospora thermoautothrophicus* comb. nov

*Charcoactinospora thermoautothrophicus* (Gr. masc. adj. *thermos,* hot; Gr. pref, *autos,* self; Gr. adj. *throphikos,* nursing, tending or feeding; N.L. masc. adj. *thermoautothrophicus,* heat-loving self-nourishing, referring to the ability to grow at high temperature at the expense of CO or H_2_ plus CO_2_.

Basonym: *Streptomyces thermoautotrophicus* Gadkari *et al.* 1990.

The description is the same as for *Streptomyces thermoautotrophicus* [31]. Phylogenomic analyses of the core genome provided strong evidence for assignment of this species in the novel genus *Charcoactinospora.* The G+C content of the type strain genome is 71 %, its approximate genome size is 5.13 Mbp, its GenBank accession is GCA_001543895.1. The type strain is UBT1^T^ (=DSM 41605^T^ = LMG 19855^T^ =NBRC 101056^T^).

### Description of *Curviacidiphilus* gen. nov

*Curviacidiphilus* (Cur.vi.a.ci.di’phi.lus L. masc. adj. *curvus,* curved or crooked; L. neut. n. *acidum,* acid; Gr. masc. adj.*philos,* loving; N.L. masc. n. *Curviacidiphilus,* curved, acidloving).

Aerobic, Gram stain positive actinobacteria, substrate hyphae are variantly colored, aerial mycelium is differentiated into long flexuous chains of spores. The pH range supporting the growth is 5.0 – 8.0. Gluconate, melibiose, D-sorbitol are used as sole carbon source at 0.1 % (w/v). L-isoleucine and L-serine are used as sole nitrogen sources at 0.1 % (w/v). Able to hydrolyse starch and chitin but unable to hydrolyse cellulose. The DNA G+C content (mol %) ranges from 71.2 to 71.8 and the genome size ranges from 8.42 to 9.54 Mbp. The genus has been separated from *Streptacidiphilus* based on 16S rRNA gene sequence and phylogenomic analyses. The type species is *Streptacidiphilus neutrinimicus* comb. nov.

### Description of *Curviacidiphilus jiangxiensis* comb. nov

*Curviacidiphilus jiangxiensis* (N.L.masc./fem.adj. *jiangxiensis*, of or pertaining to Jiangxi Province, South China, the source of the isolates)

Basonym: *Streptacidiphilus jiangxiensis* Huang *et al.* 2004.

The description is same as given before [46] for *Streptacidiphilus jiangxiensis* with the following additions. Phylogenomic analyses provided strong evidence for assignment of this species to the novel genus *Curviacidiphilus.* The G+C content of the type strain genome is 72.2 %, its approximate genome size is 9.54 Mbp, its GenBank accession is GCA_900109465.1. The type strain is 33214^T^ (=CGMCC 4.1857^T^ = DSM 45096^T^ = JCM 12277^T^ = NBRC 100920^T^).

### Description of *Curviacidiphilus pinicola* comb. nov

*Curviacidiphilus pinicola* (pi.ni.co’la L. fem. n. *pinus,* pine tree; L. masc./fem. suff. *-cola,* dweller; from L. masc./fem. n. *incola;* N.L. n. *pinicola,* referring to the habitat, pine tree soil, from which the type strain was isolated).

Basonym: *Streptacidiphilus pinicola* Roh *et al.* 2018.

The description is same as given before [47] for *Streptacidiphilus pinicola* with the following additions. Phylogenomic analyses provided strong evidence for assignment of this species to the novel genus *Curviacidiphilus.* The G+C content of the type strain genome is 71.7 %, its approximate genome size is 8.43 Mbp, its GenBank accession is GCA_003258295.1. The type strain is MMS16-CNU450^T^ (=JCM 32300^T^ = KCTC 49008^T^).

### Description of *Curviacidiphilus melanogenes* comb. nov

*Curviacidiphilus melanogenes* [me.la.no’gen.es Gr. n. *melas-anos,* black; N.L. suff. *-genes,* producing; N.L. part. adj. *melanogenes,* producing black (pigment)]

Basonym: *Streptacidiphilus melanogenes* Cho *et al.* 2008.

The description is same as given before [48] for *Streptacidiphilus melanogenes* with the following additions. Phylogenomic analyses provided strong evidence for assignment of this species to the novel genus *Curviacidiphilus.* The G+C content of the type strain genome is 71.6 %, its approximate genome size is 8.77 Mbp, its GenBank accession is GCA_000787835.1. The type strain is SB-B34^T^ (=JCM 16224^T^ = KCTC 19280^T^ = NBRC 103184^T^).

### Description of *Curviacidiphilus neutrinimicus* comb. nov

*Curviacidiphilus neutrinimicus* (neu.tri.ni.mi’cus L. adj. neut. *neutrinimicus,* here for neutral pH; L. masc. n. *inimicus,* enemy; N.L. *neutrinimicus,* enemy of the neuter (pH), nominative in opposition).

Basonym: *Streptacidiphilus neutrinimicus* Kim *et al.* 2003.

The description is same as given before [49] for *Streptacidiphilus neutrinimicus* with the following additions. Phylogenomic analyses provided strong evidence for assignment of this species to the novel genus *Curviacidiphilus.* The G+C content of the type strain genome is 71.4 %, its approximate genome size is 8.42 Mbp, its GenBank accession is GCA_000787815.1. The type strain is JL206^T^ (=DSM 41755^T^ = JCM 12365^T^ = KCTC 9911^T^ = NBRC 100921^T^).

### Description of *Curviacidiphilus anmyonensis* comb. nov

*Curviacidiphilus anmyonensis* (an.myon.en’sis N.L. masc./fem. adj. *anmyonensis,* of Anmyon, where the first strains were isolated).

Basonym: *Streptacidiphilus anmyonensis* Cho *et al.* 2008.

The description is same as given before [48] for *Streptacidiphilus anmyonensis* with the following additions. Phylogenomic analyses provided strong evidence for assignment of this species to the novel genus *Curviacidiphilus.* The G+C content of the type strain genome is 71.7 %, its approximate genome size is 9.39 Mbp, its GenBank accession is GCA_000787855.1. The type strain is AM-11^T^ (=JCM 16223^T^ = KCTC 19278^T^ = NBRC 103185^T^).

### Description of *Curviacidiphilus rugosus* comb. nov

*Curviacidiphilus rugosus* (ru.go’sus L. masc. adj. *rugosus,* wrinkled).

Basonym: *Streptacidiphilus rugosus* Cho *et al.* 2008.

The description is same as given before [48] for *Streptacidiphilus rugosus* with the following additions. Phylogenomic analyses provided strong evidence for assignment of this species to the novel genus *Curviacidiphilus.* The G+C content of the type strain genome is 71.8 %, its approximate genome size is 9.0 Mbp, its GenBank accession is GCA_000744655.1. The type strain is AM-16^T^ (=JCM 16225^T^ = KCTC 19279^T^ = NBRC 103186^T^).

### Description of *Kafeoacidophilus* gen. nov

*Kafeoacidiphilus* (Ka.feo.a.ci.di’phi.lus Gr. neut. adj. kafe, brown colour; L. neut. n. *acidum,* acid; Gr. masc. adj.*philos,* loving; N.L. adj. n. Kafeoacidiphilus, brown coloured substrate mycelium producing, acid-loving).

Aerobic, Gram positive, non-acid alcohol fast staining actinomycetes, produces brown substrate mycelium and greyish white aerial hyphae and golden-brown diffusible pigment on acidified modified Bennett’s, oatmeal and yeast extract-malt extract agar, but not on inorganic salts-starch agar. Aerial hyphae bears long flexuous chains of spores with smooth surface. Grows at 28-37 °C and at a pH of 3.0 – 6.5. D-gluconic acid, D-glucosamine hydrochloride, glycerol, *myo*inositol are used as sole carbon sources. Nitrate is reduced, but aesculin, allantoin and urea are not hydrolysed. Degrades adenine, casein, starch and uric acid, but unable to decompose elastin, guanine, hypoxanthine, Tween 80, L-tyrosine, xanthine or xylan. The genus name implies the brown coloured substrate mycelium produced by the single type species of this genus. The genus has been separated from *Streptacidiphilus* based on phylogenetic analyses of genome and 16S rRNA gene sequences. The type species is *Kafeoacidiphilus oryzae,* comb. nov.

### Description of *Kafeoacidiphilus oryzae* comb. nov

*Kafeoacidiphilus oryzae* (L. gen. fem. n. *oryzae,* of rice, denoting the isolation of the strains from a rice field).

Basonym: *Streptacidiphilus oryzae* (Wang *et al.* 2006) EMEND. Nouioui *et al.* 2018.

The description is the same as for *Streptacidiphilus oryzae* [50]. Phylogenomic analyses of the core genome provided strong evidence for assignment of this species in the novel genus *Kafeoacidiphilus.* The G+C content of the type strain genome is 73.4 %, its approximate genome size is 7.81 Mbp, its GenBank accession is GCA_000744815.1. The type strain is TH49^T^ (=CGMCC 4.2012^T^ = DSM 45098^T^ = JCM 13271^T^).

### Description of *Mangroviactinospora* gen. nov

*Mangroviactinospora* (N.L. fem. n. *mangrovi,* named for mangrove; Gr. n. *actis,* actinos a ray; Gr. fem. n. *spora,* a seed and, in biology, a spore; N.L. fem. n. *Mangroviactinospora,* mangrove actinomycete with spiny spores).

Cells stain Gram-positive and form grayish yellow aerial and substrate mycelium. Cells are positive for catalase but negative for melanoid pigment production and hemolytic activity. The cell wall peptidoglycan contained LL-diaminopimelic acid and phospholipid II type with anteiso-C_15:0_, iso-C_16:0_, iso-C_15:0_ and anteiso-C_17:0_ as predominant cellular fatty acids. Polar lipids comprised of diphosphatidylglycerol, phosphatidylinositol, phosphatidylethanolamine, hydroxyphosphatidylethanolamine, phosphatidylmethylethanolamine and hydroxyphosphatidylmethylethanolamine. The unique menaquinones identified are MK-9 (H_8_) and MK-9(H_6_). The cell wall sugars were found to be galactose, glucose, mannose, ribose and rhamnose. The type strain of the single species in the genus was isolated from mangrove forest soil at Tanjung Lumpur, Malaysia. The genomic G+C content is around 73 %. The genus has been separated from *Streptomyces* based on phylogenetic analyses of genome and 16S rRNA gene sequences. The type species is *Mangroviactinospora gilvigriseus* comb. nov.

### Description of *Mangroviactinospora gilvigriseus* comb. nov

*Mangroviactinospora gilvigriseus* (gil.vi.gri’se.us. L. adj. gilvus, yellow; L. adj. griseus, grey; N.L. masc. adj. gilvigriseus, yellow-grey, referring to the colour of the mycelium).

Basonym: *Streptomyces gilvigriseus* Ser *et al.* 2015.

The description is the same as for *Streptomyces gilvigriseus* [51]. Phylogenomic analyses of the core genome provided strong evidence for assignment of this species in the novel genus *Mangroviactinospora.* The G+C content of the type strain genome is 73 %, its approximate genome size is 5.21 Mbp, its GenBank accession is GCA_001879105.1. The type strain is MUSC 26^T^ (=DSM 42173^T^ = MCCC 1K00504^T^ = NBRC 110931^T^).

### Description of *Peterkaempfera* gen. nov

*Peterkaempfera* (N.L. masc. gen. n. *Peterkaempfera,* to honour the renowned actinomycetologist Peter Kämpfer).

An aerobic, Gram-stain-positive, non-acid-fast, non-motile, streptomycete-like actinomycete producing branched mycelium and aerial hyphae. Peptidoglycan possessed LL-diaminopimelic acid as the diagnostic diamino acid and glucose, mannose and ribose as cell-wall sugars; the major menaquinone was MK9 (H_8_, H_6_); the major fatty acids were anteiso-CA_15□: □0_ and iso-C_16□:□0_. The polar lipid profile consisted of diphosphatidylglycerol, phosphatidylethanolamine, phosphatidylinositol, glycophospholipid and aminoglycophospholipid. Utilizes cellobiose, dextrin, D-galactose, *β*-gentiobiose, D-glucose, N-acetyl-D-glucosmaine, maltose, D-mannitol, sodium bromate, trehalose and turanose as sugars. Able to grow at a pH 5 and up to 1 % (w/v) NaCl. The genomic G+C content provided in literature is around 70-73.3 mol%. The genus has been separated from *Streptacidiphilus* based on phylogenetic analyses of genome and 16S rRNA gene sequences. The type species is *Peterkaempfera griseoplanus,* comb. nov.

### Description of *Peterkaempfera bronchialis* comb. nov

*Peterkaempfera bronchialis* (bron.chi.a’lis. L. pl. n. bronchia the bronchial tubes; L. fem. suff. - alis suffix used with the sense of pertaining to; N.L. masc. adj. bronchialis pertaining to the bronchial tubes).

Basonym: *Streptacidiphilus bronchialis* Nouioui *et al.* 2019.

The description is the same as for *Streptacidiphilus bronchialis* [52]. Phylogenomic analyses of the core genome provided strong evidence for assignment of this species in the novel genus *Peterkaempfera.* The G+C content of the type strain genome is 72.7 %, its approximate genome size is 7.09 Mbp, its GenBank accession is GCA_003258605.2. The type strain is 15-057A^T^ (=DSM 106435^T^=ATCC BAA-2934^T^).

### Description of *Peterkaempfera griseoplanus* comb. nov

*Peterkampfera* griseoplanus (gri.se.o.pla’nus. L. masc. adj. griseus grey; L. masc. adj. planus flat, level; N.L. masc. adj. griseoplanus flat, grey, referring to the restricted, flat, planar growth and greyish spore colour en masse of the organism).

Basonym: *Streptacidiphilus griseoplanus* (Backus *et al.* 1957) Nouioui *et al.* 2019.

The description is the same as for *Streptacidiphilus griseoplanus* [52, 53, 54]. Phylogenomic analyses of the core genome provided strong evidence for assignment of this species in the novel genus *Peterkaempfera.* The G+C content of the type strain genome is 72.5 %, its approximate genome size is 8.25 Mbp, its GenBank accession is GCA_001418575.1. The type strain is DSM 40009^T^ (=NBRC 12779^T^=ISP 5009^T^=RIA 1046^T^=NBRC 12779^T^=CBS 505.68^T^=IFO 12779^T^=ATCC 19766^T^=AS 4.1868^T^).

### Description of *Streptantibioticus* gen. nov

*Streptantibioticus* (Gr. adj. *streptos,* pliant, twisted; N.L. masc. adj. *antibioticus,* against life, antibiotic; *Streptantibioticus).*

Aerobic, Gram positive non-acid fast staining actinomycetes. The aerial mycelium exhibits orchid pigmentation when grown on solid medium. Terminal branches of the aerial mycelium produces sporophores in compact spirals bearing spores in chains, spores are smooth, ellipsoidal to cylindrical in shape. The type strain of the single species in the genus was isolated from soil sample originating in New Jersey, U.S.A. The genus name coined owing to the different kinds of antibiotics (thienamycin, cephamycin C, penicillin N) produced by this culture and able to excrete the fluorinated antibiotic 4-fluorothreonine when cultivated in the presence of fluorine. The genomic G+C content is around 73 %. The genus has been separated from *Streptomyces* based on phylogenetic analyses of genome and 16S rRNA gene sequences. The type species is *Streptantibioticus cattleya,* comb. nov.

### Description of *Streptantibioticus cattleyae* comb. nov

*Streptantibioticus cattleyae* (N.L. gen. n. *cattleyae,* of Cattleya, a genus of orchid plants).

Basonym: *Streptomyces cattleya* Kahan *et al.* 1979.

The description is the same as for *Streptomyces cattleya* [55]. Phylogenomic analyses of the core genome provided strong evidence for assignment of this species in the novel genus *Streptantibioticus.* The G+C content of the type strain genome is 73 %, its approximate genome size is 8.10 Mbp, its GenBank accession is GCA_000240165.1. The type strain is MA4297^T^ (=ATCC 35852^T^ = DSM 46488^T^ = JCM 4925^T^ =NBRC 14057^T^ = NRRL 8057^T^).

### Emended description of *Streptomyces calvus* Backus *et al.* 1957 (Approved Lists 1980)

Heterotypic synonym: *Streptomyces asterosporus (ex* Krassilnikov 1970) Preobrazhenskaya 1986.

The species description is as given before [56] with the following additions. Spores are gray, spiny or hairy and are borne in mature spiral chains. Melanin is not produced on tyrosine agar. D-glucose, D-xylose, L-arabinose, D-rhamnose, L-rhamnose, D-galactose, raffinose, D-mannitol, I-inositol, salicin and sucrose are utilized for growth. The G+C content of the type strain genome is 72.48 % and the approximate genome size is 7.94 Mbp. The GenBank accession number for the whole-genome sequence is GCA_006782915.1. The type strain is T-3018^T^ (=AS 4.1691^T^ = ATCC 13382^T^ = ATCC 23890^T^ = BCRC 11859^T^ = CBS 350.62^T^ = CBS 676.68^T^= CCRC 11859^T^ = CECT 3271^T^ = DSM 40010^T^ = IFM 1093^T^ = IFO 13200^T^ = JCM 4326^T^ = JCM 4628^T^ = NBRC 13200^T^ = NCIMB 12240^T^ = NRRL B-2399^T^ = NRRL ISP-5010^T^ = RIA 1103^T^ = VKM Ac-1185^T^).

## Abbreviations

OGC: Orthologous gene cluster
AAI: average amino acid identity
dDDH: digital DNA-DNA hybridization
ANI: average nucleotide identity

## Acknowledgements

We also acknowledge assistance of William B. Whitman, Nikos C. Kyrpides, Tanja Woyke, Nicole Shapiro, and the other members of the JGI microbial genome sequencing team. Strain PL19 genome sequencing was conducted by the U.S. Department of Energy Joint Genome Institute, a DOE Office of Science User Facility, and was supported by the Office of Science of the U.S. Department of Energy.

## Author contribution

MM and VSS constructed the 16S phylogeny and wrote the manuscript and WSST constructed the core genome-based phylogeny and analysed the data. All authors read and approved the final version of the manuscript.

## Compliance with Ethical Standards

### Ethical approval

This article does not contain any studies with human participants or animals performed by any of the authors.

### Informed consent

Not required.

## References

1. Waksman SA, Henrici AT (1943) The nomenclature and classification of the actinomycetes. J Bacteriol 46:337–341.

2. Salam N et al (2020) Update on the classification of higher ranks in the phylum Actinobacteria. Int J Syst Evol Microbiol 70; 288–293.

3. Buchanan RE (1917) Studies in the nomenclature and classification of the *Bacteria:* II. *The primary subdivisions of the Schizomycetes*. J Bacteriol 2:155–164.

4. Kämpfer P (2015) *Streptomycetaceae* In: Whitman WB. (ed) Bergey’s Manual of Systematic of Archaea and Bacteria Wiley Online Library; pp 1–11.

5. Waksman SA, Henrici AT (1948) Family III. *Streptomycetaceae* Waksman and Henrici In: Breed RS, Murray EGD, Hitchens AP (eds), Bergey’s Manual of Determinative Bacteriology, 6^th^ ed., The Williams & Wilkins Co, Baltimore, p. 929–980.

6. Pridham TG, Tresner HD (1974) Family *Streptomycetaceae* Waksman and Henrici. In: Buchanan RE, Gibbons NE (ed.) Bergey’s manual of systematic bacteriology, 8^th^ Edn. The Williams and Wilkins, Baltimore, pp 747–748.

7. Omura S et al (1982) *Kitasatosporia*, a new genus of the order *Actinomycetales*. J Antibiot (Tokyo) 35:1013–1019.

8. Wellington EMH et al (1992) Taxonomic status of *Kitasatosporia*, and proposal unification with *Streptomyces* on the basis of phenotypic and 16S rRNA analysis and emendation of *Streptomyces* Waksman and Henrici 1943, 339AL. Int J Syst Bacteriol 42:156–160.

9. Zhang Z, Wang Y, Ruan J (1997) A proposal to revive the genus *Kitasatospora* (Omura, Takahashi, Iwai, and Tanaka 1982). Int J Syst Bacteriol 47:1048–1054.

10. Kim SB, Lonsdale J, Seong CN, Goodfellow M. *Streptacidiphilus* gen. nov., acidophilic actinomycetes with wall chemotype I and emendation of the family *Streptomycetaceae* (Waksman and Henrici (1943)AL) emend. Rainey et al. 1997. Antonie Van Leeuwenhoek 2003; 83:107–116.

11. Kämpfer P et al (2014) The family *Streptomycetaceae*. In: Rosenberg E, Delong EF, Lory S, Stackebrandt E, Thompson F. (eds) The Prokaryotes: Actinobacteria. 4^th^ Edn. Springer Reference. p.1065

12. Huang MJ et al (2017) *Allostreptomyces psammosilenae* gen. nov., sp. nov., an endophytic actinobacterium isolated from the roots of *Psammosilene tunicoides* and emended description of the family *Streptomycetaceae* [Waksman and Henrici (1943) AL] emend. Rainey et al. 1997, emend. Kim et al. 2003, emend. Zhi et al. 2009. Int J Syst Evol Microbiol 67:288–293.

13. Sahu AK et al (2017) *Allostreptomyces indica* sp. nov., isolated from India. J Antibiot (Tokyo) 70:1000–1003.

14. Nouioui I et al (2018) Genome-based taxonomic classification of the phylum *Actinobacteria*. Front Microbiol 9:2007.

15. Kämpfer P (2006) The family *Streptomycetaceae*- part 1: taxonomy. In: Dworkin M et al (eds) The Prokaryotes, vol 3, Bacteria: Firmicutes, actinomycetes. Springer, New York, pp 538–604.

16. Glaeser SP, Kämpfer P (2016) *Streptomycetaceae:* Phylogeny, Ecology and Pathogenicity. In: eLS. John Wiley & Sons, Ltd, Chichester.

17. Labeda DP et al (2012) Phylogenetic study of the species within the family *Streptomycetaceae*. Antonie Van Leeuwenhoek 101:73–104.

18. Labeda DP et al (2017) Phylogenetic relationships in the family *Streptomycetaceae* using multi-locus sequence analysis. Antonie van Leeuwenhoek 110:563–583.

19. Komaki H, Tamura T (2020) Reclassification of *Streptomyces castelarensis* and *Streptomyces sporoclivatus* as later heterotypic synonyms of *Streptomyces antimycoticus*. Int J Syst Evol Microbiol 70:288–293.

20. Chun J et al (2018) Proposed minimal standards for the use of genome data for the taxonomy of prokaryotes. Int J Syst Evol Microbiol 68: 461–466.

21. Madhaiyan M., Saravanan V.S. and See-Too W-S. (2020) Genome-based analyses reveal the presence of 12 heterotypic synonyms in the genus *Streptomyces* and emended descriptions of *Streptomyces bottropensis, Streptomyces celluloflavus, Streptomyces fulvissimus, Streptomyces glaucescens, Streptomyces murinus*, and *Streptomyces variegatus*. Int J Syst Evol Microbiol https://doi.org/10.1099/ijsem.0.004217. (in press).

22. Yoon S-H et al (2017) Introducing EzBioCloud: a taxonomically United database of 16S rRNA gene sequences and whole-genome assemblies. Int J Syst Evol Microbiol 67:1613–1617.

23. Kumar S, Stecher G, Tamura K (2016) MEGA7: Molecular Evolutionary Genetics Analysis version 7.0 for bigger datasets. Mol Biol Evol 33: 1870–1874.

24. Kimura M (1980) A simple method for estimating evolutionary rates of base substitutions through comparative studies of nucleotide sequences. J Mol Evol 16: 111–120.

25. Ding W, Baumdicker F, Neher RA (2018) panX: pan-genome analysis and exploration. Nucleic Acids Res 46: e5.

26. Darriba D et al (2020) ModelTest-NG: a new and scalable tool for the selection of DNA and protein evolutionary models. Mol Biol Evol 37: 291–294.

27. Kozlov AM et al (2019) RAxML-NG: a fast, scalable and user-friendly tool for maximum likelihood phylogenetic inference. Bioinformatics 35: 4453–4455.

28. Meier-Kolthoff JP et al (2013) Genome sequence-based species delimitation with confidence intervals and improved distance functions. BMC Bioinformatics 14: 60.

29. Blom J et al (2016) EDGAR 2.0: an enhanced software platform for comparative gene content analyses. Nucleic Acids Res 44(W1):W22–28.

30. Luo C, Rodriguez- RLM, Konstantinidis KT (2014) MyTaxa: an advanced taxonomic classifier for genomic and metagenomic sequences. Nucleic Acids Res 42:e73.

31. Gadkari D et al (1990) *Streptomyces thermoautotrophicus* sp. nov., a thermophilic CO- and H(2)-oxidizing obligate chemolithoautotroph. Appl Environ Microbiol 56:3727–3734.

32. Mackellar D et al (2016). *Streptomyces thermoautotrophicus* does not fix nitrogen. Sci Rep 6, 20086.

33. Stackebrandt E, Rainey FA, Ward-Rainey NL (1997) Proposal for a new hierarchic classification system, Actinobacteria classis nov. Int J Syst Bacteriol 47:479–491.

34. Zhi XY, Li WJ, Stackebrandt E (2009) An update of the structure and 16S rRNA gene sequence-based definition of higher ranks of the class *Actinobacteria*, with the proposal of two new suborders and four new families and emended descriptions of the existing higher taxa. Int J Syst Evol Microbiol 59:589–608.

35. Lee SD (2013) Proposal of *Motilibacteraceae* fam. nov., with the description of *Motilibacter rhizosphaerae* sp. nov. Int J Syst Evol Microbiol 63:3818–3822.

36. Shomura T et al (1983) *Actinosporangium vitaminophilum* sp. nov. Int J Syst Bacteriol 1983; 33:557–564.

37. Goodfellow M, Williams ST, Alderson G (1986) Transfer of *Actinosporangium violaceum* Krassi’nikov and Yuan, *Actinosporangium vitaminophilum* Shomura et al. and *Actinopycnidium caeruleum* Krassi’nikov to the genus *Streptomyces*, with emended descriptions of the species. Syst Appl Microbiol 8:61–64

38. Parks DH et al (2018) A standardized bacterial taxonomy based on genome phylogeny substantially revises the tree of life. Nat Biotechnol 36: 996–1004.

39. Goris J et al (2007) DNA-DNA hybridization values and their relationship to wholegenome sequence similarities. Int J Syst Evol Microbiol 57:81–91.

40. Thompson CC et al (2013) Microbial genomic taxonomy. BMC Genomics 2013; 913.

41. Stackebrandt E et al (2002) Report of the ad hoc committee for the re-evalaution of the species definition in bacteriology. Int J Syst Evol Microbiol 52: 1043–1047.

42. Liu N et al (2009) *Streptomyces alni* sp. nov., a daidzein-producing endophyte isolated from a root of *Alnus nepalensis* D. Don. Int J Syst Evol Microbiol 59:254–258.

43. Xu C et al (2006) Neutrotolerant acidophlic *Streptomyces* species isolated from acidic soils in China: *Streptomyces guanduensis* sp. nov., *Streptomyces paucisporeus* sp. nov., *Streptomyces rubidus* sp. nov. and *Streptomyces yanglinensis* sp. nov. Int J Syst Evol Microbiol 56:1109–1115.

44. Kim SB et al (2004) Taxonomic study of neutrotolerant acidophilic actinomycetes isolated from soil and description of *Streptomyces yeochonensis* sp. *nov*. Int J Syst Evol Microbiol 54:211–214.

45. Huang Y et al (2004) *Streptomyces glauciniger* sp. nov., a novel mesophilic streptomycete isolated from soil in south China. Int J Syst Evol Microbiol 54: 2085–2089

46. Huang Y et al (2004) *Streptacidiphilus jiangxiensis* sp. nov., a novel actinomycete isolated from acidic rhizosphere soil in China. Antonie Van Leeuwenhoek 86:159–165.

47. Roh SG et al (2018) *Streptacidiphilus pinicola* sp. nov., isolated from pine grove soil. Int J Syst Evol Microbiol 68:3149–3155.

48. Cho SH, Han JH, Ko HY, Kim SB (2008) *Streptacidiphilus anmyonensis* sp. nov., *Streptacidiphilus rugosus* sp. nov. and *Streptacidiphilus melanogenes* sp. nov., acidophilic actinobacteria isolated from Pinus soils. Int J Syst Evol Microbiol 58:1566–1570.

49. Kim SB et al (2004) Taxonomic study of neutrotolerant acidophilic actinomycetes isolated from soil and description of *Streptomyces yeochonensis* sp. *nov*. Int J Syst Evol Microbiol 54: 211–214.

50. Wang L et al (2006) *Streptacidiphilus oryzae* sp. nov., an actinomycete isolated from rice-field soil in Thailand. Int J Syst Evol Microbiol 56:1257–1261.

51. Ser HL et al (2015) *Streptomyces gilvigriseus* sp. nov., a novel actinobacterium isolated from mangrove forest soil. Antonie Van Leeuwenhoek 107: 1369–1378.

52. Nouioui I et al. (2019) *Streptacidiphilus bronchialis* sp. nov., aciprofloxacin-resistant bacterium from a human clinical specimen; reclassification of *Streptomyces griseoplanus* as *Streptacidiphilus griseoplanus* comb. nov. and emended description of the genus *Streptacidiphilus*. Int J Syst Evol Microbiol 69:1047–1056

53. Backus EJ, Tresner HD, Campbell TH (1957) The nucleocidin and alazopetin producing organisms: two new species of *Streptomyces*. Antibiot Chemother (Northfield) 7:532–541.

54. Kämpfer P. (2012) Genus incertae sedis II. *Streptacidiphilus* Kim, Lonsdale, Seong and Goodfellow 2003a, 1219VP (Effective publication: Kim, Lonsdale, Seong and Goodfellow 2003b, 115.). In: Goodfellow M et al (editors). Bergey’s Manual of Systematic Bacteriology, 2nd ed, vol. 5. The Actinobacteria, Part B. New York: Springer; pp. 1777–1805.

55. Kahan JS et al (1979) Thienamycin, a new β-lactam antibiotic discovery, taxonomy, isolation and physical properties. J Antibiot (Tokyo) 30: 1–12.

56. Backus EJ, Tresner HD, Campbell TH (1957) The nucleocidin and alazopetin producing organisms: two new species of *Streptomyces*. Antibiot Chemother (Northfield) 7:532–541.

57. Kim SB et al. *Streptacidiphilus* gen. nov., acidophilic actinomycetes with wall chemotype I and emendation of the family *Streptomycetaceae* (Waksman and Henrici (1943)AL) emend. Rainey et al. 1997. Antonie Van Leeuwenhoek 83:107–116.

58. Shirling EB, Gottlieb D (1977) Retrospective evaluation of International *Streptomyces* Project taxonomic criteria. In Actinomycetes: the Boundary Microorganisms (edited by Arai). University Park Press, Baltimore, pp. 9–41.

59. Ōmura S, Takahashi Y, Iwai Y (1989) Genus *Kitasatosporia*. In: Williams, Sharpe and Holt (ed.) Bergey’s Manual of Systematic Bacteriology, vol. 4, Williams & Wilkins, Baltimore, pp. 2594–2598.

60. Lonsdale JT (1985) Aspects of the biology of acidophilic actinomycetes. PhD thesis, University of Newcastle, Newcastle upon Tyne.

61. Williams ST, Goodfellow M, Alderson G (1989) Genus *Streptomyces* Waksman and Henrici. In: Williams, Sharpe and Holt (ed.) Bergey’s Manual of Systematic Bacteriology, vol. 4. Williams & Wilkins, Baltimore, pp. 2452–2492.

62. Nakagaito Y et al. Proposal of *Streptomyces atroaurantiacus* sp. nov. and *Streptomyces kifunensis* sp. nov. and transferring *Kitasatosporia cystarginea* Kusakabe and Isono to the genus *Streptomyces* as *Streptomyces cystargineus* comb. nov. J Gen Appl Microbiol 38:627–633.

63. Antony-Babu S, Goodfellow M (2008) Biosystematics of alkaliphilic streptomycetes isolated from seven locations across a beach and dune sand system. Antonie van Leeuwenhoek 94: 581–591

64. Komaki H et al. (2020) Diversity of PKS and NRPS gene clusters between *Streptomyces abyssomicinicus* sp. *nov. and its taxonomic neighbor*. J Antibiot (Tokyo) 73:141–151.

65. Komaki H, Tamura T (2019) Reclassification of *Streptomyces rimosus* subsp. Paromomycinus as *Streptomyces paromomycinus* sp. nov. Int J Sys Evol Microbiol 69: 2577–2583.

66. Huang MJ et al. (2017) *Allostreptomyces psammosilenae* gen. nov., sp. nov., an endophytic actinobacterium isolated from the roots of *Psammosilene tunicoides* and emended description of the family *Streptomycetaceae* [Waksman and Henrici (1943) AL] emend. Rainey et al. 1997, emend. Kim et al. 2003, emend. Zhi et al. 2009. Int J Syst Evol Microbiol 67: 288–293.

67. Ping X et al. (2004) *Streptomyces scabrisporus* sp. nov. Int J Syst Evol Microbiol 54: 577–581.

68. Nagai A et al. (2011) *Streptomyces aomiensis* sp. nov., isolated from a soil sample using the membrane-filter method. Int J Syst Evol Microbiol 61:947–950.

69. Mikami Y, Miyashita K, Arai T (1982) Diaminopimelic acid profilesof alkalophilic and alkaline-resistant strains of Actinomycetes. J Gen Microbiol 128: 1709–1712.

70. Kroppenstedt RM (1985) Fatty acid and menaquinone analysis of actinomycetes and related organisms. In: Goodfellow and Minnikin (ed.) Chemical Methods in Bacterial Systematics, Academic Press, London, pp. 173–199.

