## Supplementary Material for "Genomic and phylogenomic insights into the family *Streptomycetaceae* lead to proposal of *Charcoactinosporaceae* fam. nov. and 8 novel genera with emended descriptions of *Streptomyces calvus*"

**Table S3.** List of the core genes in the genome used for phylogenomic analysis.

| NCBI Protein Accession | Gene |
|---|---|
| WP_074993204.1 | NUDIX hydrolase |
| WP_070028582.1 | YggS family pyridoxal phosphate-dependent enzyme |
| WP_074992763.1 | ParB/RepB/Spo0J family partition protein |
| WP_070022023.1 | lipoyl(octanoyl) transferase LipB |
| WP_070025151.1 | FABP family protein |
| WP_070027039.1 | heat-inducible transcriptional repressor HrcA |
| WP_074992865.1 | folate-binding protein YgfZ |
| WP_074992658.1 | recombination protein RecR |
| WP_074991826.1 | HIT domain-containing protein |
| WP_070024163.1 | adenylosuccinate synthase |
| WP_009190566.1 | anti-sigma regulatory factor |
| WP_071828679.1 | preprotein translocase subunit SecG |
| WP_070026304.1 | 50S ribosomal protein L13 |
| WP_009190144.1 | 30S ribosomal protein S5 |
| WP_014674378.1 | 30S ribosomal protein S8 |
| WP_070026314.1 | 50S ribosomal protein L5 |
| WP_009300593.1 | 30S ribosomal protein S13 |
| WP_003998809.1 | 50S ribosomal protein L36 |
| WP_070026864.1 | WhiB family transcriptional regulator |
| WP_023550764.1 | 50S ribosomal protein L27 |
| WP_070027822.1 | ATP synthase F0 subcomplex C subunit |
| WP_039656223.1 | 30S ribosomal protein S17 |
| WP_070024148.1 | DUF5063 domain-containing protein |
| WP_009190136.1 | 50S ribosomal protein L16 |
| WP_070024021.1 | MoaD/ThiS family protein |
| WP_070028831.1 | 30S ribosomal protein S6 |
| WP_003956500.1 | 50S ribosomal protein L34 |
| WP_070025179.1 | glycine--tRNA ligase |
| WP_018544333.1 | DUF3117 domain-containing protein |
| WP_003974250.1 | 50S ribosomal protein L30 |
| WP_070022292.1 | 30S ribosomal protein S16 |
| WP_070023573.1 | 2-phospho-L-lactate transferase |
| WP_074993184.1 | PHP domain-containing protein |

**Table S4.** Similarity in polyphasic data observed between *S. asterosporus* DSM 41452$^T$ and *S. calvus* CECT 3271$^T$

| **Characteristic†** | ***Streptomyces asterosporus* DSM 41452$^T$** | ***Streptomyces calvus* CECT 3271$^T$** |
|---|---|---|
| Morphological characters: | | |
| Spore color | Gray | Gray |
| Spore chain morphology | Spira | Spira |
| Melanoid pigment production | – | – |
| Spore wall ornamentation | Spiny | Hairy |
| Physiological tests: | | |
| D-Glucose | + | + |
| D-Xylose | − | + |
| L-Arbinose | + | + |
| D-Rhamnose | + | + |
| L-Rhamnose | + | + |
| D-Galactose | + | + |
| Raffinose | − | + |
| D-Mannitol | + | + |
| i-Inositol | n | + |
| Salicin | n | + |
| Sucrose | n | + |
| G+C content (mol%) | 72.5 | 72.5 |
| Genome size (Mbp) | 7.77 | 7.94 |

+, positive; −, negative; n-not determined.

†Data from Kämpfer et al. (2014).

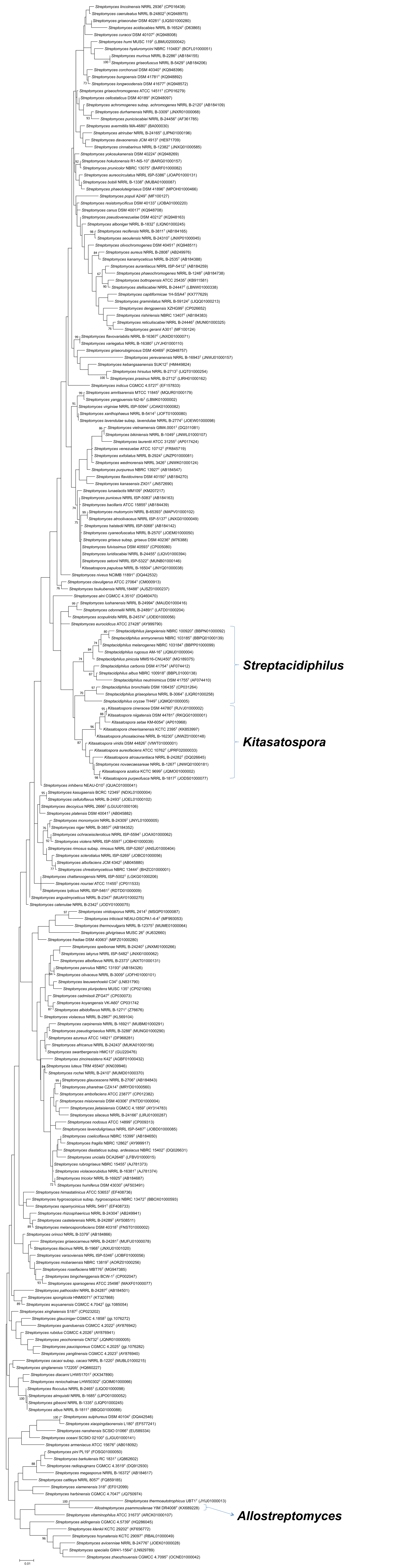
Streptacidiphilus
Kitasatospora
Allostreptomyces
0.01

**Fig. S1.** Maximum likelihood tree based on 16S rRNA gene sequences showing the phylogenetic relationship among *Streptomyces* and closely related genera belongs to *Streptomycetaceae* family members. The numbers at the nodes are given as percent and represent the levels of bootstrap support (>70%) based on maximum likelihood analyses of 500 re-sampled data sets. Scale bar, 0.01 nt substitutions per position.

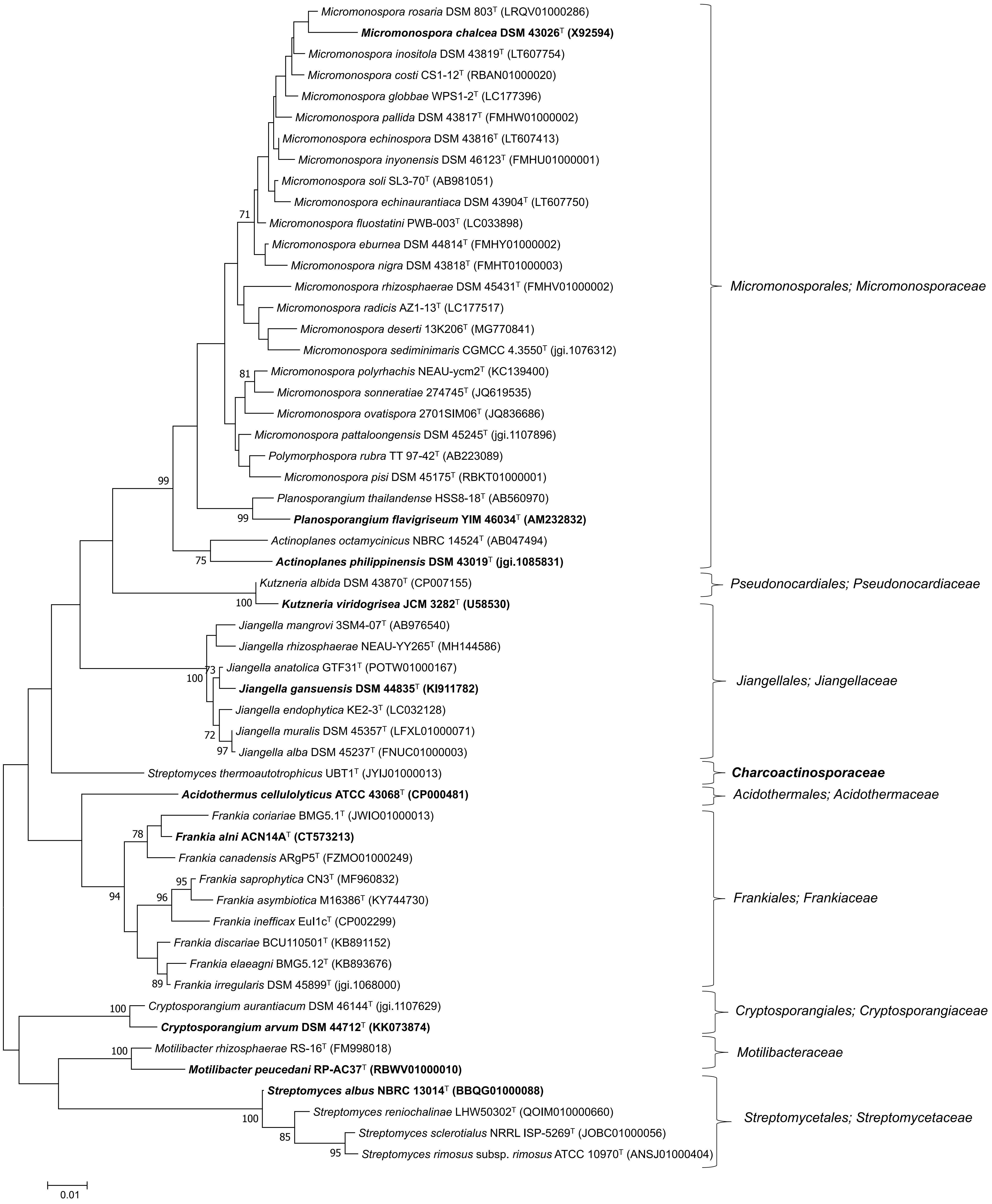

Micromonospora rosaria DSM 803$^{T}$ (LRQV01000286)
**Micromonospora chalcea DSM 43026$^{T}$ (X92594)**
Micromonospora inositola DSM 43819$^{T}$ (LT607754)
Micromonospora costi CS1-12$^{T}$ (RBAN01000020)
Micromonospora globbae WPS1-2$^{T}$ (LC177396)
Micromonospora pallida DSM 43817$^{T}$ (FMHW01000002)
Micromonospora echinospora DSM 43816$^{T}$ (LT607413)
Micromonospora inyonensis DSM 46123$^{T}$ (FMHU01000001)
Micromonospora soli SL3-70$^{T}$ (AB981051)
Micromonospora echinaurantiaca DSM 43904$^{T}$ (LT607750)
71
Micromonospora fluostatini PWB-003$^{T}$ (LC033898)
Micromonospora eburnea DSM 44814$^{T}$ (FMHY01000002)
Micromonospora nigra DSM 43818$^{T}$ (FMHT01000003)
Micromonospora rhizosphaerae DSM 45431$^{T}$ (FMHV01000002)
Micromonospora radicis AZ1-13$^{T}$ (LC177517)
Micromonospora deserti 13K206$^{T}$ (MG770841)
Micromonospora sediminimaris CGMCC 4.3550$^{T}$ (jgi.1076312)
81
Micromonospora polyrhachis NEAU-ycm2$^{T}$ (KC139400)
Micromonospora sonneratiae 274745$^{T}$ (JQ619535)
Micromonospora ovatispora 2701SIM06$^{T}$ (JQ836686)
Micromonospora pattaloongensis DSM 45245$^{T}$ (jgi.1107896)
Polymorphospora rubra TT 97-42$^{T}$ (AB223089)
Micromonospora pisi DSM 45175$^{T}$ (RBKT01000001)
99
Planosporangium thailandense HSS8-18$^{T}$ (AB560970)
99
**Planosporangium flavigriseum YIM 46034$^{T}$ (AM232832)**
Actinoplanes octamycinicus NBRC 14524$^{T}$ (AB047494)
75
**Actinoplanes philippinensis DSM 43019$^{T}$ (jgi.1085831)**
Micromonosporales; Micromonosporaceae
Kutzneria albida DSM 43870$^{T}$ (CP007155)
100
**Kutzneria viridogrisea JCM 3282$^{T}$ (U58530)**
Pseudonocardiales; Pseudonocardiaceae
Jiangella mangrovi 3SM4-07$^{T}$ (AB976540)
Jiangella rhizosphaerae NEAU-YY265$^{T}$ (MH144586)
73
Jiangella anatolica GTF31$^{T}$ (POTW01000167)
100
**Jiangella gansuensis DSM 44835$^{T}$ (KI911782)**
Jiangella endophytica KE2-3$^{T}$ (LC032128)
72
Jiangella muralis DSM 45357$^{T}$ (LFXL01000071)
97
Jiangella alba DSM 45237$^{T}$ (FNUC01000003)
Jiangellales; Jiangellaceae
Streptomyces thermoautotrophicus UBT1$^{T}$ (JYIJ01000013)
**Charcoactinosporaceae**
**Acidothermus cellulolyticus ATCC 43068$^{T}$ (CP000481)**
Acidothermales; Acidothermaceae
Frankia coriariae BMG5.1$^{T}$ (JWIO01000013)
78
**Frankia alni ACN14A$^{T}$ (CT573213)**
Frankia canadensis ARgP5$^{T}$ (FZMO01000249)
95
Frankia saprophytica CN3$^{T}$ (MF960832)
94
96
Frankia asymbiotica M16386$^{T}$ (KY744730)
Frankia inefficax EuI1c$^{T}$ (CP002299)
Frankia discariae BCU110501$^{T}$ (KB891152)
Frankia elaeagni BMG5.12$^{T}$ (KB893676)
89
Frankia irregularis DSM 45899$^{T}$ (jgi.1068000)
Frankiales; Frankiaceae
100
Cryptosporangium aurantiacum DSM 46144$^{T}$ (jgi.1107629)
**Cryptosporangium arvum DSM 44712$^{T}$ (KK073874)**
Cryptosporangiales; Cryptosporangiaceae
100
Motilibacter rhizosphaerae RS-16$^{T}$ (FM998018)
**Motilibacter peucedani RP-AC37$^{T}$ (RBWV01000010)**
Motilibacteraceae
**Streptomyces albus NBRC 13014$^{T}$ (BBQG01000088)**
100
Streptomyces reniochalinae LHW50302$^{T}$ (QOIM010000660)
85
Streptomyces sclerotialus NRRL ISP-5269$^{T}$ (JOBC01000056)
95
Streptomyces rimosus subsp. rimosus ATCC 10970$^{T}$ (ANSJ01000404)
Streptomycetales; Streptomycetaceae
0.01

**Fig. S2.** Maximum likelihood tree of the class *Actinobacteria* families showing the presence of 9 orders/families based on 16S rRNA gene sequence comparison. The phylogenetic relatedness of the families of the class *Actinobacteria* is outlined. Taxa newly described in this paper are highlighted in bold. The numbers at the nodes are given as percent and represent the levels of bootstrap support (>70%) based on maximum likelihood analyses of 1000 re-sampled data sets. Scale bar, 0.01 nt substitutions per position.

**Supplementary references**

**Kämpfer P, Glaeser SP, Parks L, van Keulen G, Dyson P.** The family *Streptomycetaceae*. In: Rosenberg E, Delong EF, Lory S, Stackebrandt E, Thompson F. (editors) *The Prokaryotes: Actinobacteria*. 4th Edn. Springer Reference. 2014; p.1065

**Tajima K, Takahashi Y, Seino A, Iwai Y, Omura S.** Description of two novel species of the genus *Kitasatospora* Omura et al. 1982, *Kitasatospora cineracea* sp. nov. and *Kitasatospora niigatensis* sp. nov. *Int J Syst Evol Microbiol* 2001; 51:1765-1771.
